# Paradoxical SERCA dysregulation contributes to atrial fibrillation in a model of diet-induced obesity

**DOI:** 10.1101/2024.08.02.606385

**Authors:** Daniela Ponce-Balbuena, Daniel J. Tyrrell, Carlos Cruz-Cortés, Guadalupe Guerrero-Serna, Andre Monteiro Da Rocha, Todd J. Herron, Jianrui Song, Danyal S. Raza, Justus Anumonwo, Daniel R. Goldstein, L. Michel Espinoza-Fonseca

**Author notes:** Correspondence to: L. Michel Espinoza-Fonseca. These authors contributed equally (Daniela Ponce-Balbuena, Daniel J. Tyrrell, Carlos Cruz-Cortés, Guadalupe Guerrero-Serna, Andre Monteiro Da Rocha). Present address: Department of Medicine, University of Wisconsin, Madison, WI, USA. Present address: Department of Molecular & Cellular Pathology, The University of Alabama at Birmingham, Birmingham, AL, USA. **Competing Interest Statement:** D.P.-B., D.J.T., C.C.-C., G.G.S., A.M.D.R., J.S., D.S.R., J.A., D.R.G., and L.M.E.-F. declare no competing interests. T.J.H is co-founder of Cartox, Inc., and scientific advisor to StemBioSys, Inc. Author contribution statement: Conceptualization: J.A., D.R.G. and L.M.E.-F. Investigation: D.P.-B., D.J.T., C.C.-C., G.G.S., A.M.D.R., J.S., D.S.R. and J.A. Formal Analysis: D.P.-B., D.J.T., C.C.-C., G.G.S., A.M.D.R., T.J.H., J.A., D.R.G. and L.M.E.-F. analyzed data Supervision: J.A., D.R.G., and L.M.E.-F. Funding acquisition: D.R.G. and L.M.E.-F. Writing and draft editing: D.R.G. and L.M.E.-F.

## Abstract

Obesity is a major risk factor for atrial fibrillation (AF) the most common serious cardiac arrhythmia, but the molecular mechanisms underlying diet-induced AF remain unclear. In this study, we subjected mice to a chronic high-fat diet and acute sympathetic activation (‘two-hit’ model) to study the mechanisms by which diet-induced obesity promotes AF. Surface electrocardiography revealed that diet-induced obesity and sympathetic activation synergize during intracardiac tachypacing to induce AF. At the cellular level, diet-induced obesity and acute adrenergic stimulation facilitate the formation of delayed afterdepolarizations in atrial myocytes, implicating altered Ca^2+^ dynamics as the underlying cause of AF. We found that diet-induced obesity does not alter the expression of major Ca^2+^-handling proteins in atria, including the sarcoplasmic reticulum Ca^2+^-ATPase (SERCA), a major component of beat-to-beat Ca^2+^ cycling in the heart. Paradoxically, obesity reduces phospholamban phosphorylation, suggesting decreased SERCA activity, yet atrial myocytes from obese mice showed a significantly increased Ca^2+^ transient amplitude and SERCA-mediated Ca^2+^ uptake. Adrenergic stimulation further increases the Ca^2+^ transient amplitude but does not affect Ca^2+^ reuptake in atrial myocytes from obese mice. Transcriptomics analysis showed that a high-fat diet prompts upregulation of neuronatin, a protein that has been implicated in obesity and is known to stimulate SERCA activity. We propose a mechanism in which obesity primes SERCA for paradoxical activation, and adrenergic stimulation facilitates AF conversion through a Ca^2+^-induced Ca^2+^ release gain in atrial myocytes. Overall, this study links obesity, altered Ca^2+^ signaling, and AF, and targeting this mechanism may prove effective for treating obesity-induced AF.

## Introduction

In the last several decades, the rate of obesity has progressively increased and is now one of the leading causes of morbidity and mortality in the world. According to the World Health Organization, 1 in 8 people in the world are currently living with obesity; 2.2 billion adults were overweight in 2022 and 890 million were considered obese [1]. It is estimated that by 2030 about half of the world’s population will suffer from obesity, and about 11% of individuals will be morbidly obese [2]. Obesity contributes to cardiovascular risk factors, including dyslipidemia, diabetes, hypertension, and sleep disorders. Obesity also leads to the development of cardiovascular disease and cardiovascular disease mortality independently of other cardiovascular risk factors [3].

Diet-induced obesity is a major risk factor for atrial fibrillation (AF) the most common serious cardiac arrhythmia in the developed world [4–6]. AF represents the leading cause of hospitalization, affecting nearly five million patients across the United States [7]. There is evidence indicating that obese individuals have ∼50% increased risk compared to non-obese patients [8]. Other studies have shown that obese young men have more than a 2-fold risk of AF compared with young men of normal weight [8]. Clinical studies have associated increased body mass index with AF, where every unit increase in body mass index (BMI) increases the development of AF by up to 8% [5, 9]. While obesity occurs with other risk factors for AF, including hypertension, atherosclerosis, diabetes, and sleep apnea, a recent study showed demonstrated that obesity alone promotes AF [10]. Moreover, obesity is associated with a higher recurrence of AF [11]. Inflammation and remodeling of ion transport have been shown to contribute to obesity-induced AF [12, 13], yet the pathophysiological and molecular mechanisms associated with AF are complex and the molecular basis for this association remains unclear. Therefore, there is an urgent need to investigate the mechanisms by which obesity increases the risk for AF, ultimately revealing new pathways that can lead to therapeutic interventions for this condition.

In this study, we used a ‘two-hit’ mouse model (chronic high-fat diet and acute sympathetic activation) to study the molecular mechanisms by which diet-induced obesity promotes AF. We found that diet-induced obesity and sympathetic activation synergize during intracardiac tachypacing to induce AF. Diet-induced obesity and acute adrenergic stimulation facilitate the formation of delayed afterdepolarizations in atrial myocytes, suggesting that altered Ca^2+^ dynamics are the underlying mechanism promoting AF in obese mice. Obesity does not affect the expression of major Ca^2+^-handling proteins, including the sarcoplasmic reticulum Ca^2+^-ATPase (SERCA), a major component of beat-to-beat Ca^2+^ cycling in the heart. Paradoxically, diet-induced obesity decreases phospholamban phosphorylation, yet atrial myocytes from obese mice showed a significantly increased SERCA-mediated Ca^2+^ uptake. Adrenergic stimulation further increases the Ca^2+^ transient amplitude but does not affect Ca^2+^ reuptake in atrial myocytes from obese mice. A high-fat diet upregulates neuronatin, a protein that has been implicated in obesity and is known to stimulate SERCA activity. These findings suggest a novel mechanism where obesity primes SERCA for activation, altering Ca^2+^ signaling in atrial myocytes and leading to atrial fibrillation during sympathetic activation.

## Results

### Diet-induced obesity induces metabolic and inflammatory imbalances without fibrosis or alterations in cardiovascular hemodynamics

We used an established obesity model in which mice are fed a diet high in fat for 8 weeks [14] to examine the effect of obesity on the induction of AF. We used a group of mice fed a regular chow diet as a control. Mice fed a high-fat diet gained substantial weight and fat mass (Fig. 1A,B) but maintained a similar lean mass over the 8-week feeding period compared to the mice fed a regular diet (Fig. 1C). A high-fat diet produced both glucose intolerance and insulin resistance (Fig. 2A,B), in agreement with previous studies linking a high-fat diet with metabolic remodeling [15]. We also found that diet-induced obesity increased the secretion of tumor necrosis factor-*α* (TNF-*α*), interleukin-6 (IL-6), and galectin-3 (Fig. 2C). These findings agree with previous studies showing that obesity increases the levels of TNF-*α* and IL-6 [16, 17].

**Fig. 1.**
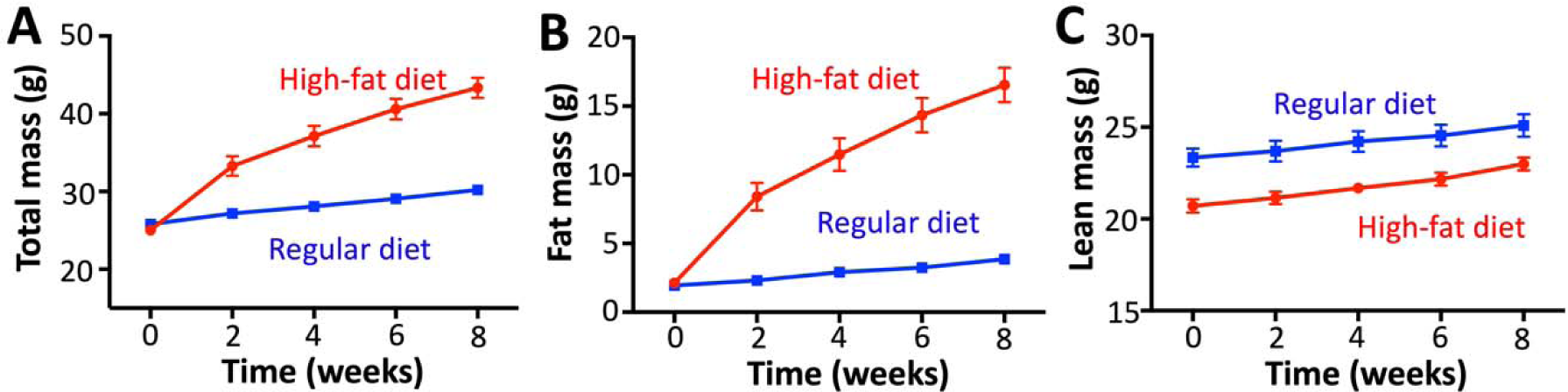
Changes in body weight and composition induced by a high-fat diet. We monitored (A) body mass, (B) fat mass, and (C) lean mass of mice fed regular and high-fat diets for eight weeks (N=15 mice per group).

**Fig. 2.**
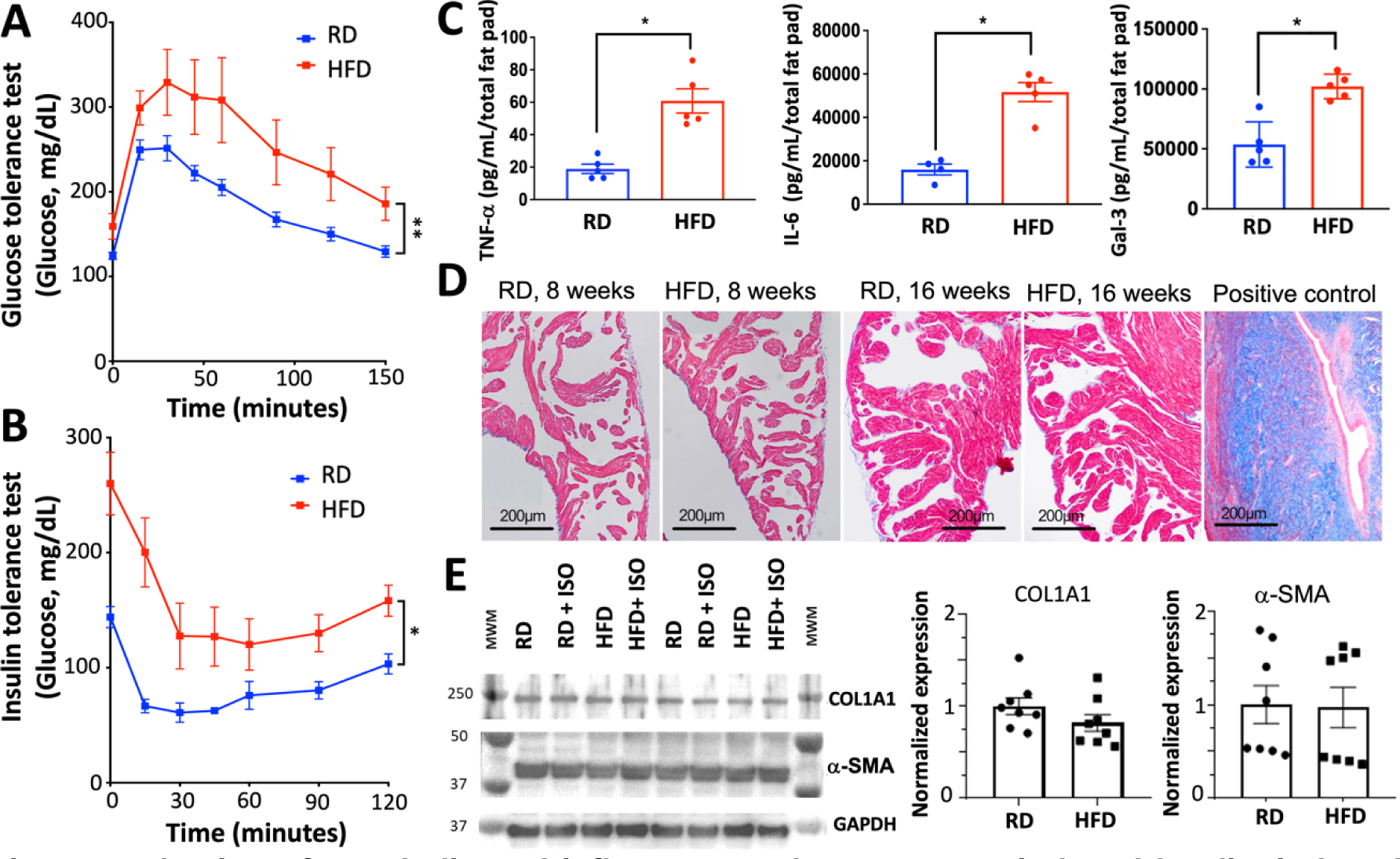
**Evaluation of metabolic and inflammatory derangements induced by diet-induced obesity in mice**. (A) Glucose tolerance test of mice fed a regular high-fat diet for 8 weeks; N=4 mice fed a regular diet, N=5 mice fed a high-fat diet. (B) Insulin tolerance test of mice fed a regular and high-fat diet for 8 weeks; N=3 mice fed a regular diet, N=4 mice fed a high-fat diet. (C) ELISA-based quantification of tumor necrosis factor α (TNFα), interleukin 6 (IL-6), and galectin-3 (Gal-3) in the gonadal white adipose tissue of mice fed regular and high-fat diets; N=4 mice for each group. (D) Masson’s trichrome staining of atria of mice fed regular and high-fat diets for 8 weeks or 16 weeks; for comparison, we show fibrosis in rat uterus as a positive control. (E) Western blot analysis of atria of mice fed either regular or high-fat diets for 8 weeks with or without acute isoproterenol (ISO) treatment. Western blot analysis of COL1A1 and α-SMA was normalized against GAPDH in all atrial tissue samples. N=8 mice per group. RD, regular diet; HFD, high-fat diet. \**p*<0.05, \*\**p*<0.01, two-tailed t-test

A high-fat diet prompted the expected metabolic and inflammatory alterations in mice, but histological examination did not show evidence of fibrosis in the atria of mice fed a high-fat diet (Fig. 2D). Extending the diet for 16 weeks did not induce fibrosis (Fig. 2D), thus the duration of the diet is not a factor for the lack of fibrosis in our model. Western blot analysis did not show significant changes in the expression of pro-fibrotic proteins in the atria of obese mice, including collagen and α-smooth muscle actin (α-SMA) (Fig. 2E). Echocardiography studies showed that a high-fat diet does not alter heart rate, left ventricle ejection fraction, left ventricular mass, or left atrial dimensions (Table 1). Non-invasive blood pressure assessment at the end of the diet showed that there are no significant differences in either diastolic or systolic blood pressure between mice fed a high-fat diet versus a regular chow diet (Table 1). Collectively, these findings indicate that mice fed a high-fat diet undergo the expected inflammatory and metabolic imbalances that are typical of diet-induced obesity, but without apparent fibrosis or alterations in cardiovascular hemodynamics.

**Table 1.**
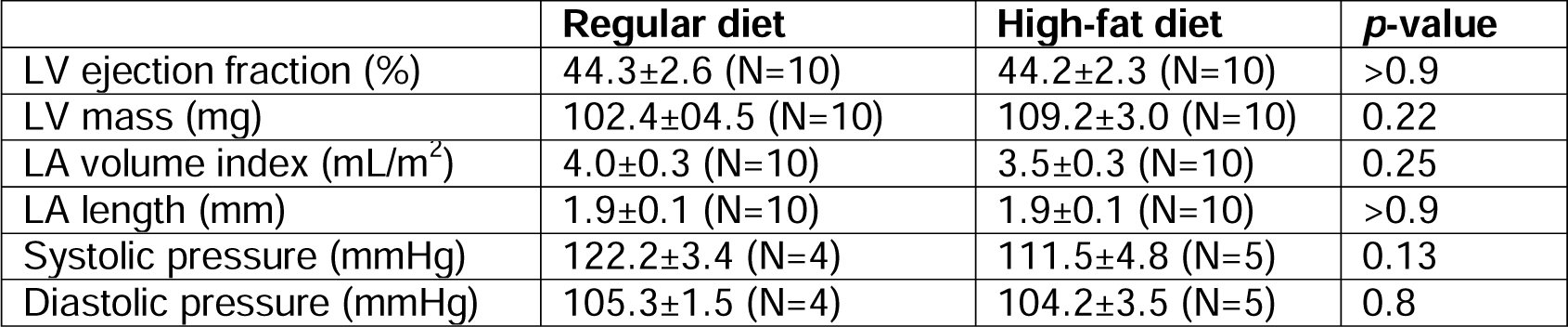
Functional cardiac parameters for mice fed regular and high-fat diets.

### Diet-induced obesity and acute sympathetic activation synergize during intracardiac tachypacing to induce atrial fibrillation

We used surface electrocardiogram (ECG) analysis to determine whether a high-fat diet alone is sufficient to increase susceptibility to atrial arrhythmias before applying intracardiac electrical stimulation. AF was determined as the occurrence of rapid and fragmented atrial electrograms (lack of regular P-waves) with irregular ventricular rhythm (irregular RR intervals) lasting at least 3 seconds [18]. A representative intracardiac recording of AF is shown in Fig. 3A. Following an 8- week feeding period, the P-wave duration, PR interval, and QRS duration were similar between mice fed high-fat and regular diets (Table 2). However, mice fed a high-fat diet presented prolonged RR interval duration (Table 2). Alterations in the autonomic nervous system can predispose to arrhythmia, including AF [19, 20], so we studied the effects of acute sympathetic activation. Upon isoproterenol administration, but before applying intracardiac electrical stimulation, mice fed a high-fat diet exhibited a significantly shorter RR interval duration compared to mice fed a regular diet (Table 3). However, obese mice did not have changes in P-wave duration, PR interval, and QRS duration (Table 3).

**Fig. 3.**
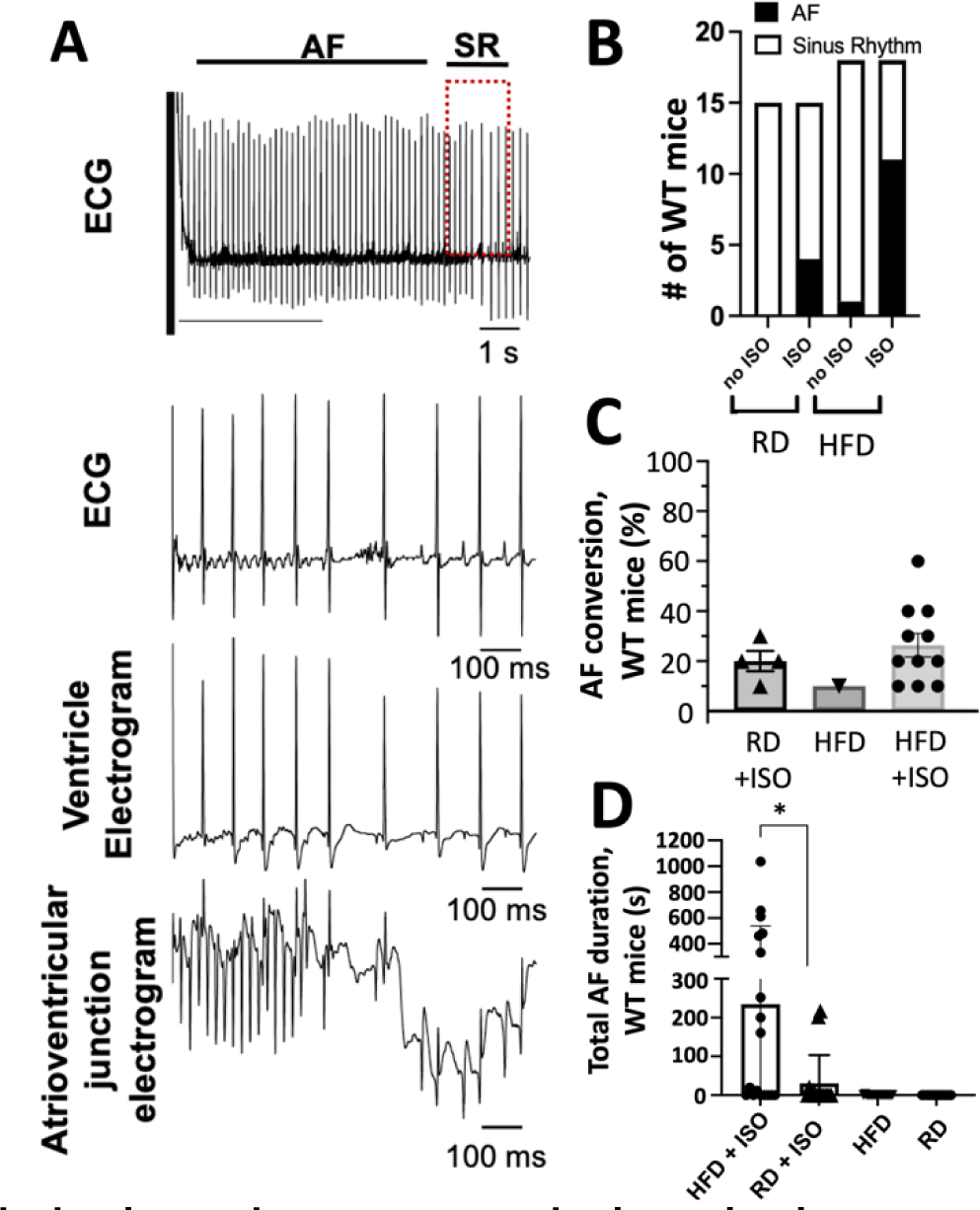
**Diet-induced obesity and acute sympathetic activation synergize to induce atrial fibrillation in mice**. (A) Representative recordings of lead II surface ECG with simultaneous ventricular and atrioventricular junction intracardiac electrograms. We show an event of AF induced by obesity in mice after the heart was paced, AF stopped spontaneously and was followed by a normal sinus rhythm (SR). Expanded signals show that the AF event spontaneously stops, followed by a normal SR (red box) in a mouse fed a high-fat diet. (B) The number of mice within each treatment group that exhibited AF (black area) or remained in sinus rhythm (white area). (C) The number of AF conversions; mice were given 10 tachypacing attempts to convert into AF. (D) Mice that were fed a high-fat diet and acutely administered isoproterenol (ISO) exhibited significantly longer AF episodes compared to mice fed a regular diet and treated acutely with ISO. RD, regular diet; HFD, high-fat diet. \**p*<0.05, two-tailed t-test.

**Table 2.**
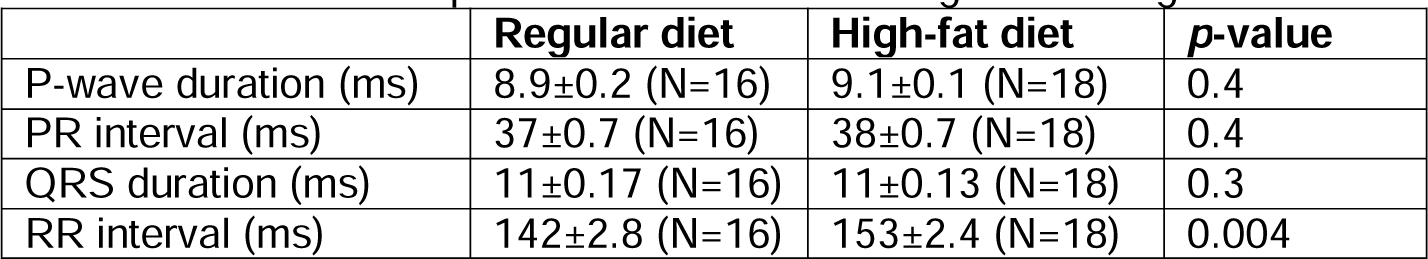
Surface ECG parameters for mice fed regular and high-fat diets.

**Table 3.** Surface ECG parameters before and after isoproterenol treatment of mice fed regular and high-fat diets.

|  | **Regular diet, basal (N=10)** | **High-fat diet, basal (N=9)** | **p-value** | **Regular diet, ISO (N=10)** | **High-fat diet, ISO (N=9)** | **p-value** |
| --- | --- | --- | --- | --- | --- | --- |
| P-wave duration (ms) | 8.5±0.4 | 8.6±0.4 | 0.8 | 8.1±0.6 | 8.8±0.3 | 0.3 |
| PR interval (ms) | 37±1.7 | 35±0.9 | 0.3 | 35±1.8 | 30±1.6 | 0.07 |
| QRS duration (ms) | 11±0.4 | 10±0.1 | 0.1 | 11±0.5 | 10±0.1 | 0.1 |
| RR interval (ms) | 124.5±3.7 | 125.6±3.1 | 0.8 | 102±1.3 | 91±1.4 | <0.0001 |

Alterations in the RR interval duration are a significant biomarker for AF detection [21]. Therefore, we used intracardiac electrical stimulation to determine the rate of conversion of atrial fibrillation induced by diet-induced obesity. Mice fed high-fat and regular diets were subjected to tachypacing in the right atrium to induce AF. Upon intracardiac electrical stimulation and in the absence of isoproterenol, mice fed a standard diet did not convert to AF (Fig. 3B). Following tachypacing and isoproterenol administration, 26% of mice on a regular diet converted to AF (Fig. 3B); however, this difference was not significant compared to mice fed a regular diet but without isoproterenol treatment (*p*=0.09; Fig. 3B). Conversely, 61% of mice on a high-fat diet treated acutely with isoproterenol developed AF following tachypacing; this represents a significant increase compared to obese mice without isoproterenol administration (*p*=0.0009). We determined the ease of conversion and duration of AF by performing ten tachypacing attempts on mice fed both high-fat and regular diets that exhibited AF events in the presence or absence of isoproterenol treatment. Eleven mice that were fed a high-fat diet and treated with isoproterenol exhibited an increased ease of AF conversion compared to mice fed a regular diet that converted to AF after treatment without isoproterenol (Fig. 3C). Furthermore, mice that were fed a high-fat diet exhibited a significantly longer AF duration compared to mice fed a regular diet (*p*=0.02; Fig. 3D). In the absence of isoproterenol, mice fed high-fat and regular diets did not exhibit AF events for more than 10 seconds (Fig. 3D). Collectively, these findings indicate that diet-induced obesity and acute sympathetic activation synergize during intracardiac tachypacing to induce atrial fibrillation.

### Effects of diet-induced obesity on the action potential characteristics of atrial myocytes

We performed patch-clamp electrophysiology studies to analyze the action potential (AP) characteristics of single isolated atrial myocytes from mice fed a high-fat diet and compared them with isolated atrial myocytes from animals fed a regular diet. APs were analyzed at 1-Hz stimulation; representative recordings from myocytes from the left and right atria are shown in Fig. 4A. A high-fat diet does not induce significant differences in the action potential amplitude, overshoot, resting membrane potential (RPM), or maximum upstroke velocity (dV/dt_max_) in myocytes isolated from either left or right atrium (Fig. 4B). We also analyzed the effects of diet-induced obesity on the action potential duration (APD) at 25% (APD_25_), 50% (APD_50_), and 90% (APD_90_) depolarization. Compared to a regular diet, a high-fat diet does not induce significant changes in APD_25_, APD_50,_ or APD_90_ of myocytes isolated from the right atrium (Fig. 4C). However, a high-fat diet significantly increased the APD_25_ by ∼4 ms *vs* a regular diet (*p*=0.04) and APD_50_ by

**Fig. 4.**
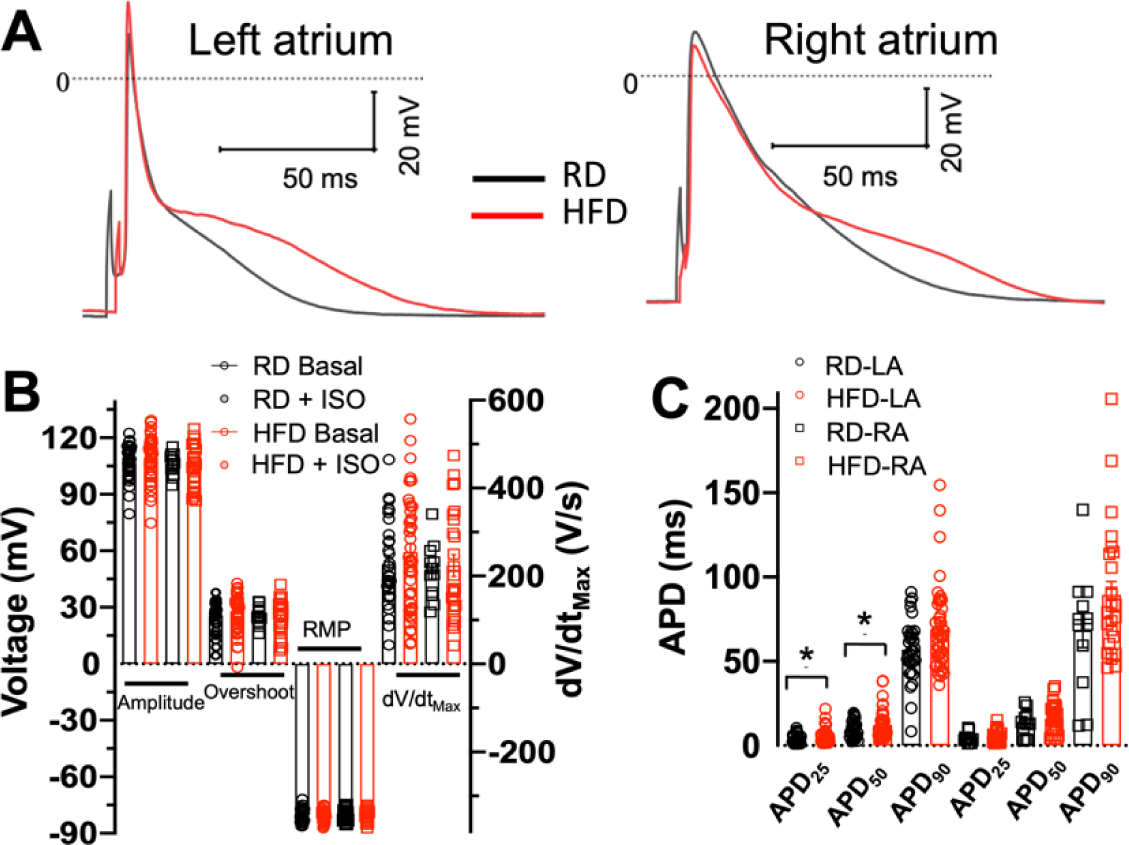
Effects of diet-induced obesity on the action potential characteristics of atrial myocytes. (A) Representative basal action potentials from left and right atrial myocytes isolated from mice fed regular and high-fat diets at 1-Hz stimulation. (B) Amplitude, overshoot, resting membrane potential (RMP), and maximum upstroke velocity (dV/dt_max_) of myocytes from mice fed regular (black) and high-fat (red) diets in the presence and absence of isoproterenol (ISO) treatment. (C) Action potential duration (APD) to 25, 50, and 90% of repolarization. For these experiments, we used the left atrium of 6 animals, and the right atrium of 5 animals fed a regular diet; we used the left and right atria of 5 mice fed a high-fat diet. RD, regular diet; HFD, high-fat diet; LA, left atrium; RA, right atrium. For mice fed a regular diet, we used N=6 mice, n=12 cells from the left atrium, and N=5 animals, n=12 cells from the right atrium. For mice fed a high-fat diet, we used N=5 animals, n=12 cells from the left atrium, and N=5 mice, n=25 cells from the right atrium. \**p*<0.05, two-tailed t-test.

∼8 ms *vs* a regular diet (*p*=0.03) in cardiomyocytes isolated from the left atrium (Fig. 4C).

We next evaluated the effects of isoproterenol on AP characteristics in left atrial myocytes. Representative recordings are shown in Fig. 5A. Isoproterenol does not induce significant differences in the action potential amplitude, overshoot, RPM, or dV/dt_max_ from baseline recordings (i.e., before isoproterenol treatment) in both diet groups (Fig 5B). Isoproterenol prolongs the APD_25_ by ∼30% (4.8±0.6 ms to 6.4±0.7 ms) and the APD_50 by_ ∼18% (11±1.3 ms to 13±1 ms) of left atrium myocytes isolated from mice fed a regular diet. While a high-fat diet prolongs both APD_25_ and APD_50_, it also impairs the response of atrial cells to *β*-adrenergic stimulation. This impaired response to isoproterenol is illustrated in Fig. 5C, showing a significant difference in the change in APD from baseline upon treatment with isoproterenol. These findings indicate that a high-fat diet significantly prolongs the action potential of left atrial myocytes and impairs the effects of *β*-adrenergic stimulation on APD.

**Fig. 5.**
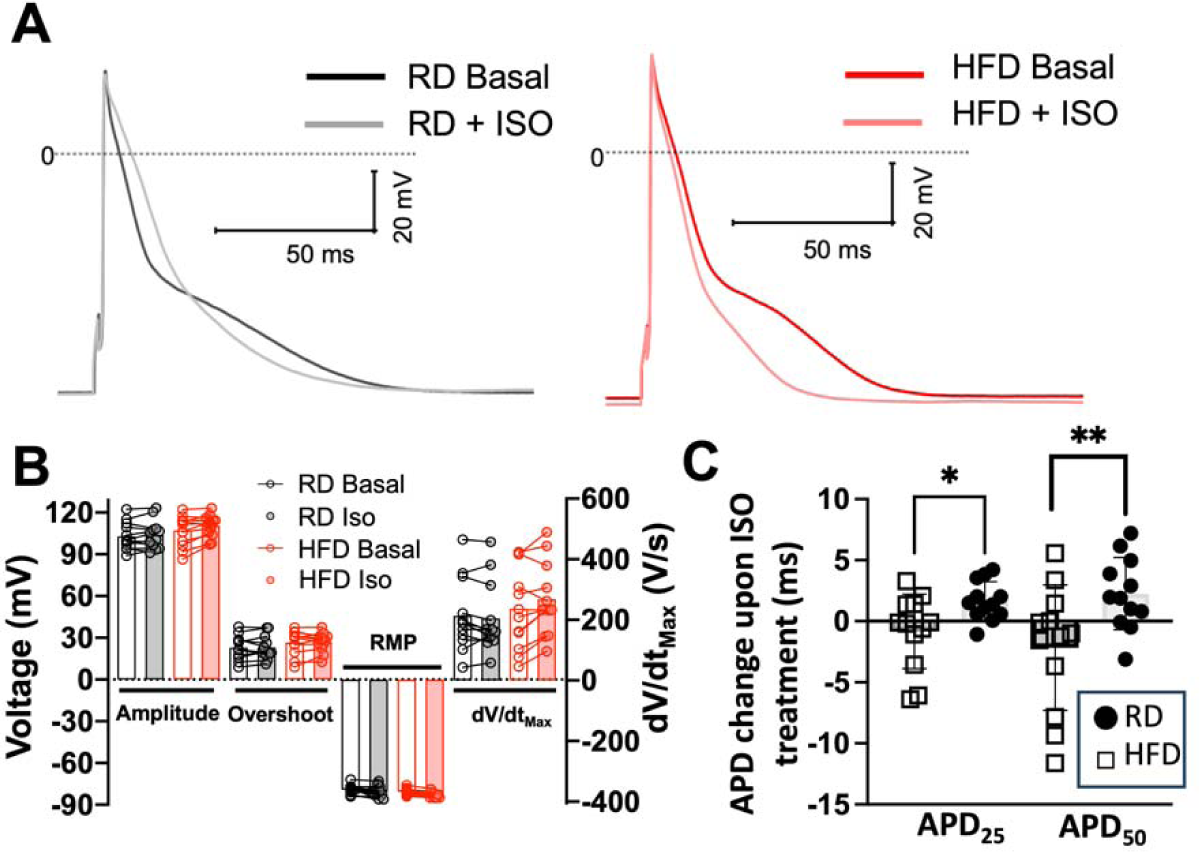
Effects of diet-induced obesity on the action potential characteristics of left atrial myocytes upon isoproterenol treatment. (A) Representative basal action potentials from left atrial myocytes isolated from mice fed regular and high-fat diets at 1-Hz stimulation; measurements are shown at basal and isoproterenol (ISO) treatment conditions. (B) Amplitude, overshoot, resting membrane potential (RMP), and maximum upstroke velocity (dV/dt_max_) of myocytes isolated from left atria. (C) Changes in APD_25_ and APD_50_ in response to isoproterenol (ISO) in mice fed regular and high-fat diets. RD, regular diet; HFD, high-fat diet. N=6 mice, n=12 cells; \**p*<0.05, \*\**p*<0.01, two-tailed t-test.

### Diet-induced obesity does not alter I_Ca_ and I_K_ densities but produces delayed afterdepolarization events in atrial myocytes

The changes in AP observed in left atrial myocytes from mice fed a high-fat diet can be produced by alterations in the balance between depolarizing inward calcium (I_Ca_) and outward repolarizing potassium (I_K_) currents [22]. Therefore, we measured I_Ca_ and I_K_ densities in atrial myocytes before and after isoproterenol treatment. Cell capacitance values were 53±6 pF for myocytes isolated from mice fed a regular diet, and 53±2 pF for myocytes isolated from mice fed a high-fat diet; no statistically significant differences were found between both groups (*p*=0.9). Voltage-clamp studies did not show differences in I_Ca_ and I_K_ at basal conditions in atrial myocytes from mice fed a regular *vs* high-fat diet (Fig. 6). Peak I_Ca_ values at 0 mV were 3.9±0.5 pA/pF (regular diet) and - 3.6±0.2 pA/pF (high-fat diet), with no significant differences found between groups (*p*=0.5; Fig. 6A,B). Peak I_K_ values at 40 mV were 16±1.5 pA/pF (regular diet) *vs* 14±0.7 pA/pF (p=0.29; Fig. 6C,D). Overall, a high-fat diet does not induce alterations in inward depolarizing I_Ca_ or total I_K_ outward repolarizing currents in atrial myocytes.

**Fig. 6.**
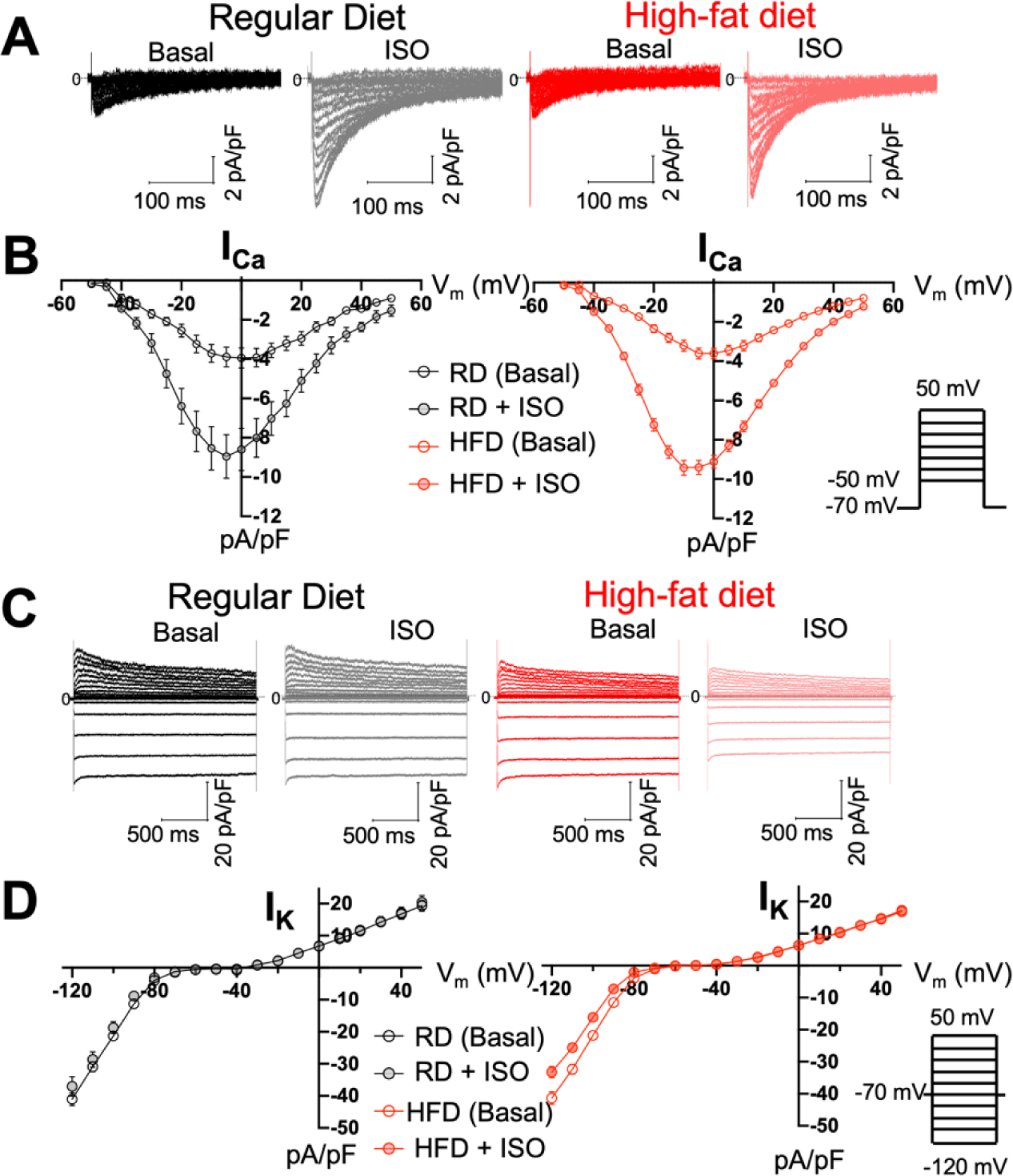
**Diet-induced obesity does not induce changes in inward Ca^2+^ and outward K^+^ currents in atrial myocytes**. (A) Representative I_Ca_ traces from left atrial myocytes from mice fed regular and high-fat diets at baseline (basal) and upon isoproterenol treatment. (B) I_Ca_ current- voltage (I/V) relationship at baseline (basal) and under adrenergic stimulation; we used N=3 mice, n=6 cells from mice fed a regular diet, N=3 mice, n=10 cells from obese mice. (C) Representative I_K_ traces from left atrial myocytes from mice fed regular and high-fat diets at baseline (basal) and after isoproterenol treatment. (D) I_K_ current-voltage (I/V) relationship before and after isoproterenol treatment; we used N=3 mice, n=8 cells from mice fed a regular diet, N=3 mice, n=10 cells from obese mice. The inserts in panels B and D show the voltage/pulse protocols that were applied to record the total Ca^2+^ and K^+^ currents, respectively. RD, regular diet; HFD, high-fat diet.

We performed current-clamp experiments to determine whether a high-fat diet produces delayed afterdepolarizations (DADs) in atrial myocytes. The production of DADs was measured at 2-Hz stimulation [23, 24]. Representative AP traces showed that atrial myocytes from mice fed a regular diet do not show DADs in the presence or absence of isoproterenol (Fig. 7A). While a high-fat diet alone is sufficient to produce a significant increase of DADs in atrial myocytes (Fig. 7B,C), isoproterenol increases the formation of DADs by ∼150% (Fig. 7C). These findings indicate that a high-fat diet produces DADs in atrial myocytes, and that isoproterenol increases the incidence of DADs induced by diet-induced obesity in mice.

**Fig. 7.**
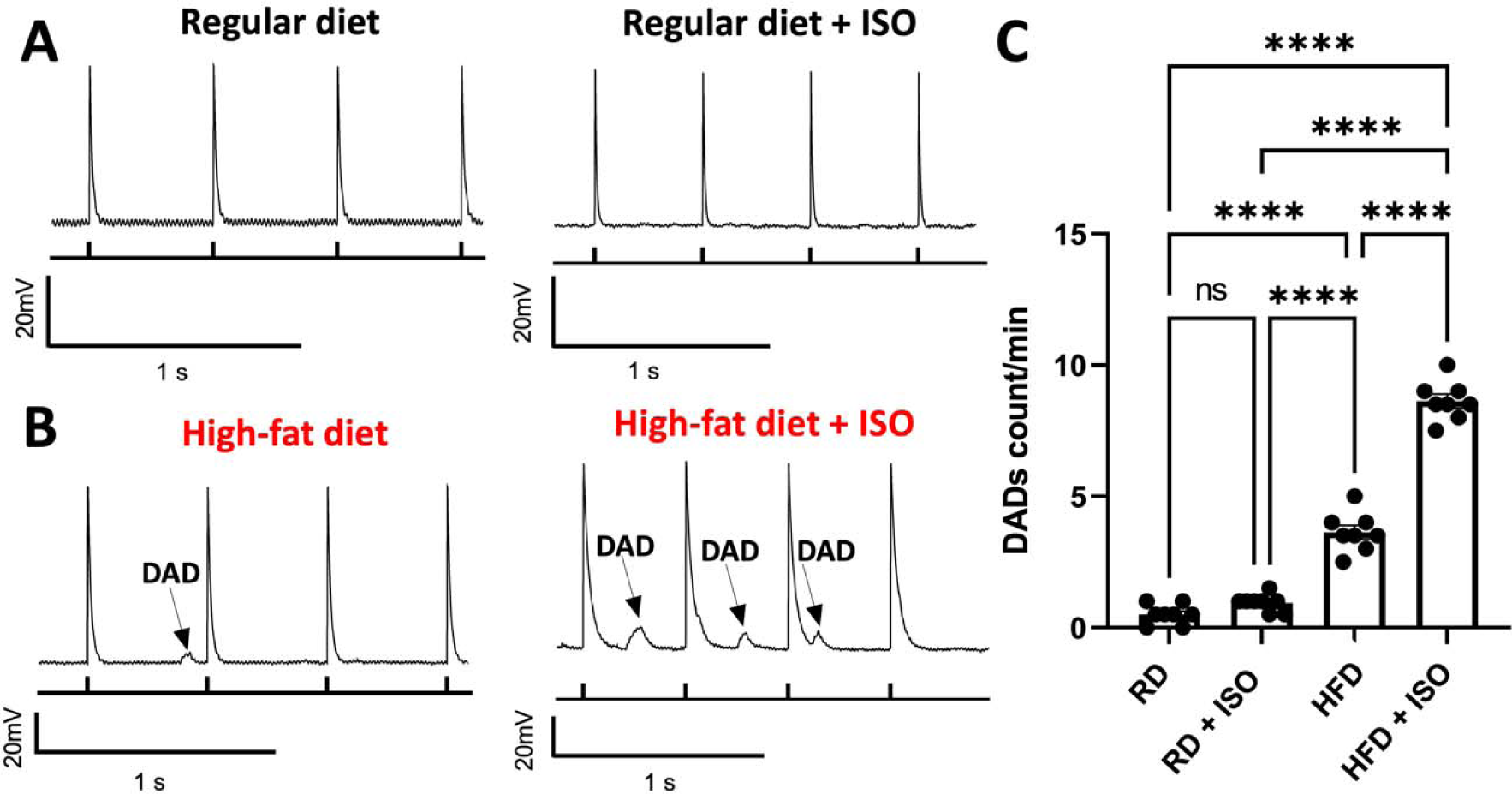
**DAD incidence is enhanced in atrial myocytes isolated from obese mice.** (A) Representative recordings of atrial myocytes isolated from mice fed (A) a regular diet and (B) a high-fat diet. Recordings are shown in the absence and presence of isoproterenol. (C) Quantification of the DAD formation events. Under current clamp conditions, cells were paced at a frequency of 2Hz. The incidence of DADs was analyzed over a 2-minute recording period and compared across the experimental groups. RD, regular diet; HFD, high-fat diet. RD, regular diet; HFD, high-fat diet. For each group, we recorded DADs from N=5 mice and n=8 cells. Analysis was performed using ANOVA with Tukey’s post-hoc test; \*\*\*\**p*<0.0001.

### Diet-induced obesity induces paradoxical dysregulation of SERCA2a in atrial myocytes

A hallmark of DADs is the dysregulation of intracellular Ca^2+^ dynamics in cardiac cells [25]. Therefore, we used Western blot analysis to determine if diet-induced obesity induces changes in the expression of proteins within the atria that are involved in intracellular Ca^2+^ dynamics. Compared to a regular diet, a high-fat diet did not induce significant changes in the expression of the ryanodine receptor (RyR), the L-type Ca^2+^ channel (Ca_v_1.2), the cardiac Na^+^-Ca^2+^ exchanger (NCX1), and the cardiac sarcoplasmic reticulum Ca^2+^-ATPase (SERCA2a) (Fig. 8A,B). We also determined changes in phospholamban (PLN) expression and phosphorylation. PLN exists as monomers that directly inhibit SERCA2a activity, and as oligomers that serve either as a storage or effector of SERCA2a activity. Compared to mice fed a regular diet, there is a significant increase in the presence of monomeric, but not oligomeric form of PLN in atria of mice fed a high-fat diet (Fig. 8A,B). Atria from mice fed a high-fat diet showed significantly lower basal (i.e., before isoproterenol treatment) PLN phosphorylation and no response to isoproterenol treatment compared to atria from mice fed a regular diet (Fig. 8C,D). Our findings indicate that diet-induced obesity impairs PLN phosphorylation in the atria of mice fed a high-fat diet.

**Fig. 8.**
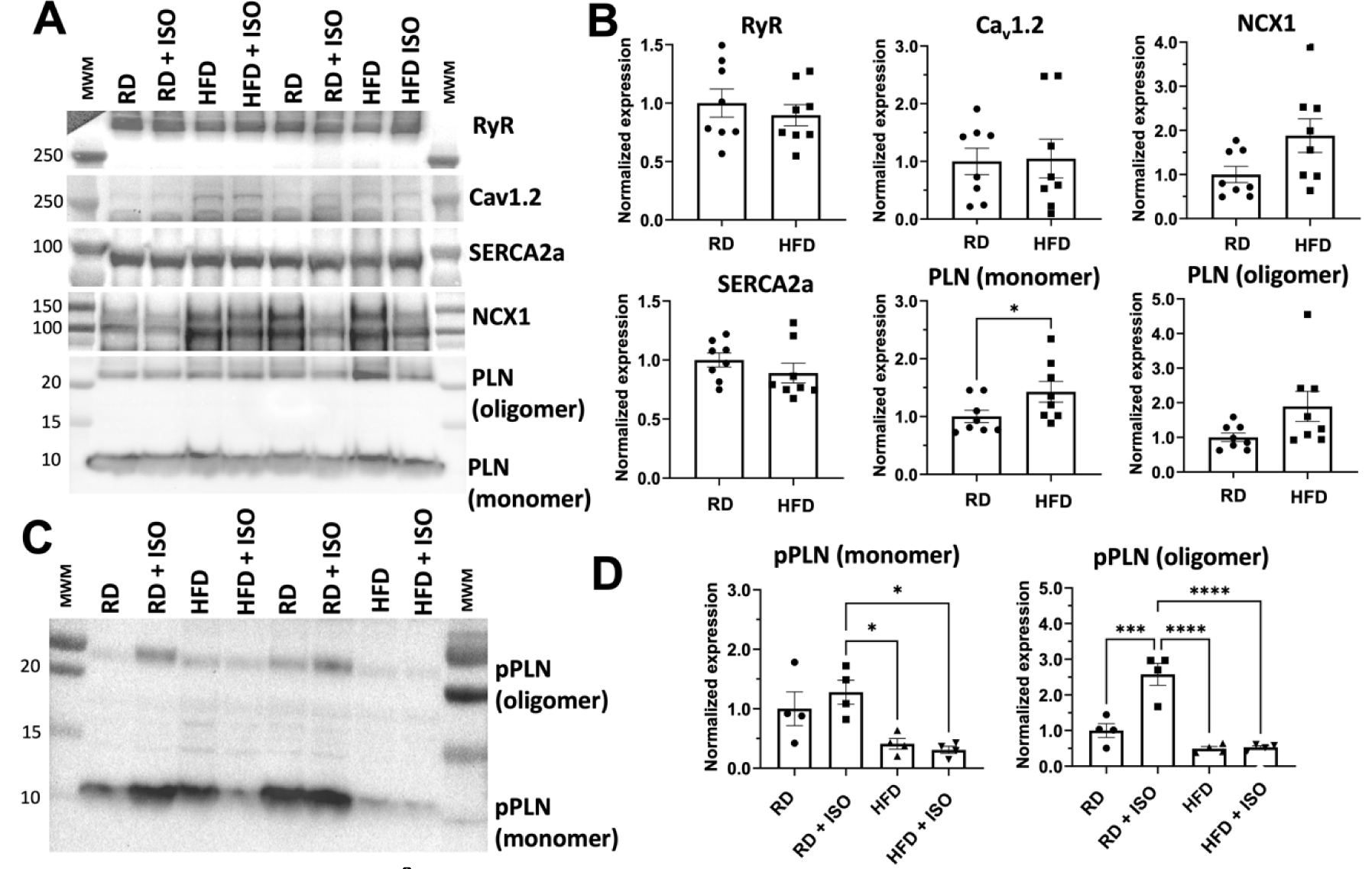
Expression of Ca^2+^-handling proteins in atrial tissue induced by diet-induced obesity. (A) Western blot analysis of proteins involved in intracellular Ca^2+^ cycling in atria, including the ryanodine receptor (RyR), the L-type calcium channel (Ca_v_1.2), SERCA2a, the Na^+^/Ca^2+^ exchanger (NCX1) and PLN. (B) Quantification of protein expression by Western blot analysis of the Ca^2+^ handling proteins shown in panel A. Protein expression was normalized against GAPDH in all atrial tissue samples; N=8 mice. (C) Western blotting of PLN phosphorylation in atrial tissue of mice fed regular and high-fat diets, with and without isoproterenol (ISO) treatment. (D) Protein quantification of phosphorylated phospholamban in atrial lysates; N=4 mice. RD, regular diet; HFD, high-fat diet. Statistical differences were tested using ANOVA with Tukey post-hoc test \**p* < 0.05, *** *p* < 0.001, **** *p* < 0.0001.

It has been shown that impaired PLN phosphorylation inhibits SERCA2a and reduces Ca^2+^ reuptake into the sarcoplasmic reticulum [26]. Therefore, we used single-cell Ca^2+^ imaging to establish the effects of a high-fat diet on intracellular Ca^2+^ dynamics in atrial myocytes. Ca^2+^ transients were analyzed at 1-Hz and 2-Hz field stimulation. The stimulated Ca^2+^ transient amplitude was significantly increased in myocytes from mice fed a high-fat diet compared to those from mice on a regular diet (Fig. 9A). Unexpectedly, the Ca^2+^ transient decay, *τ*, is significantly faster in atrial myocytes from mice fed a high-fat diet *vs* a regular diet (Fig. 9B). Treatment of myocytes with the SERCA2a inhibitor thapsigargin significantly slows the Ca^2+^ transient decay in myocytes from mice fed a high-fat diet, e.g., average *τ* values of ∼100 ms and ∼450 ms before and after thapsigargin treatment at 2-Hz stimulation (*p*<0.0001). This indicates that the effect on the Ca^2+^ transient decay found here is largely mediated by SERCA2a activation. Isoproterenol treatment of atrial myocytes from mice fed a high-fat diet significantly increased Ca^2+^ transient amplitude (relative to baseline) compared to myocytes from mice fed a regular diet (Fig. 9C). Isoproterenol treatment does not affect the relative Ca^2+^ transient decay of myocytes from obese *vs* nonobese mice (Fig. 9D).

**Fig. 9.**
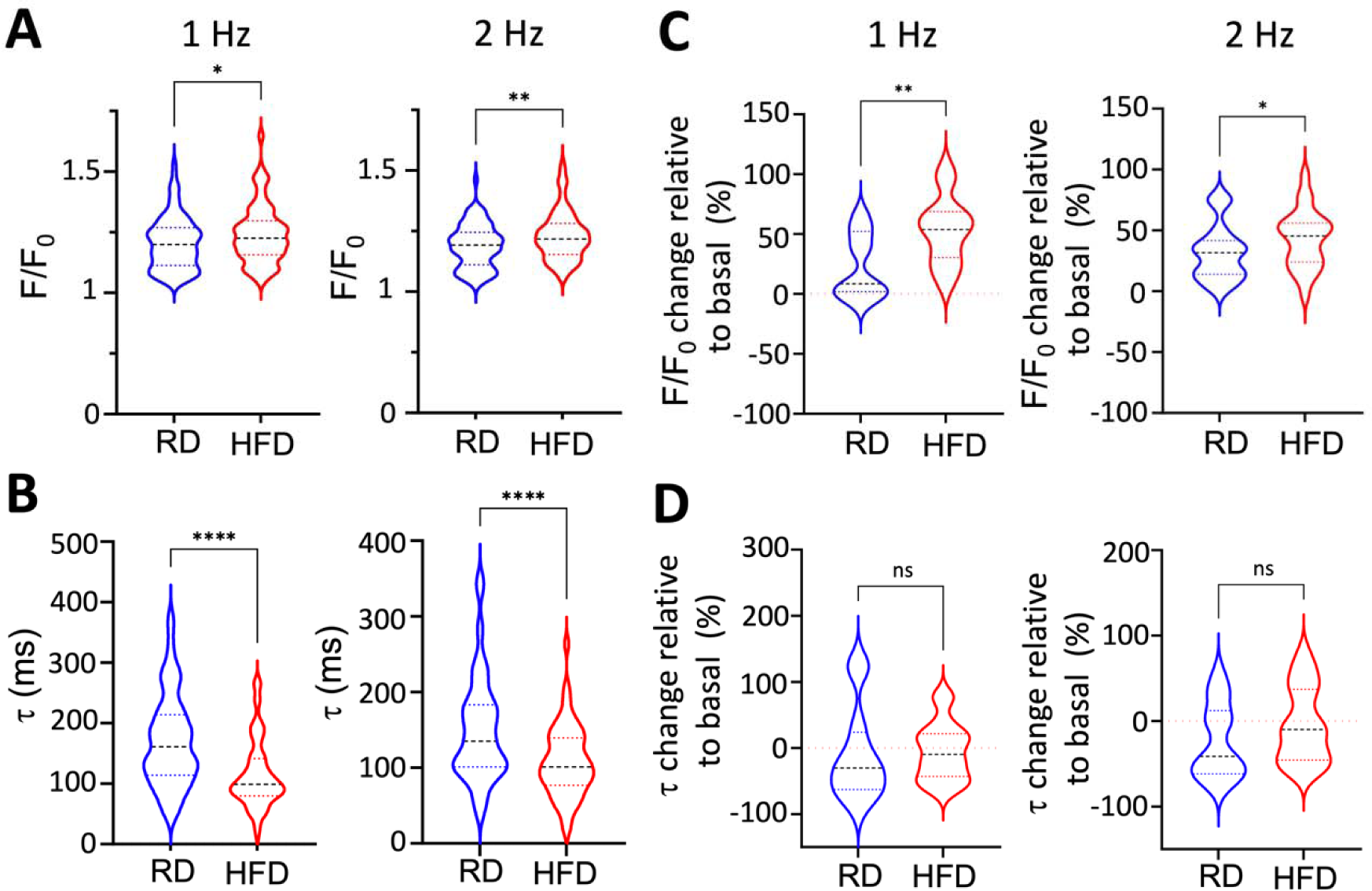
**Effects of a high-fat diet on the intracellular calcium transient in atrial myocytes**. (A) Ca^2+^ transient amplitude and (B) Ca^2+^ transient decay (*τ*) measured in atrial myocytes isolated from mice fed regular and high-fat diets at 1- and 2-Hz stimulation. Changes in (C) Ca^2+^ transient amplitude and (D) Ca^2+^ transient decay (*τ*) in response to isoproterenol for each group (regular and high-fat diets) at 1- and 2-Hz stimulation. Changes in these parameters are relative to the basal signal, i.e., before isoproterenol treatment. Data are presented as a violin plot, where dashed lines represent quartiles, full lines represent the median, and widths represent the number of individuals with the same value of the measured parameter. RD, regular diet; HFD, high-fat diet. N=9 mice fed a regular diet, N=8 mice fed a high-fat diet. Statistical differences were tested using the Mann-Whitney U-test; \**p* < 0.05, ** *p* < 0.01, **** *p* < 0.0001.

### Diet-induced atrial upregulates genes involved in metabolism and stress

We performed RNA sequencing to obtain the transcriptomic signature of atrial tissue in our model of diet-induced obesity. The analysis was performed on atrial tissue from five mice fed a high-fat diet, and five mice fed a regular diet as a control. After adjusting for known covariates and correcting for multiple comparisons, we found 83 differentially expressed genes between mice fed a high-fat diet and control (FDR <0.05, Fig. 10A,B). Of these genes, 24 were upregulated and 59 were downregulated (Fig. 10B and Tables S1 and S2 of the Supplementary Information). Gene ontology pathway analysis of the differentially expressed genes showed that a high-fat diet primarily affects the expression of genes involved in atrial metabolic processes including organic acid metabolism, lipid metabolism, lipid catabolism, fatty acid oxidation, and small-molecule catabolic processing (Fig. 10C).

**Fig. 10.**
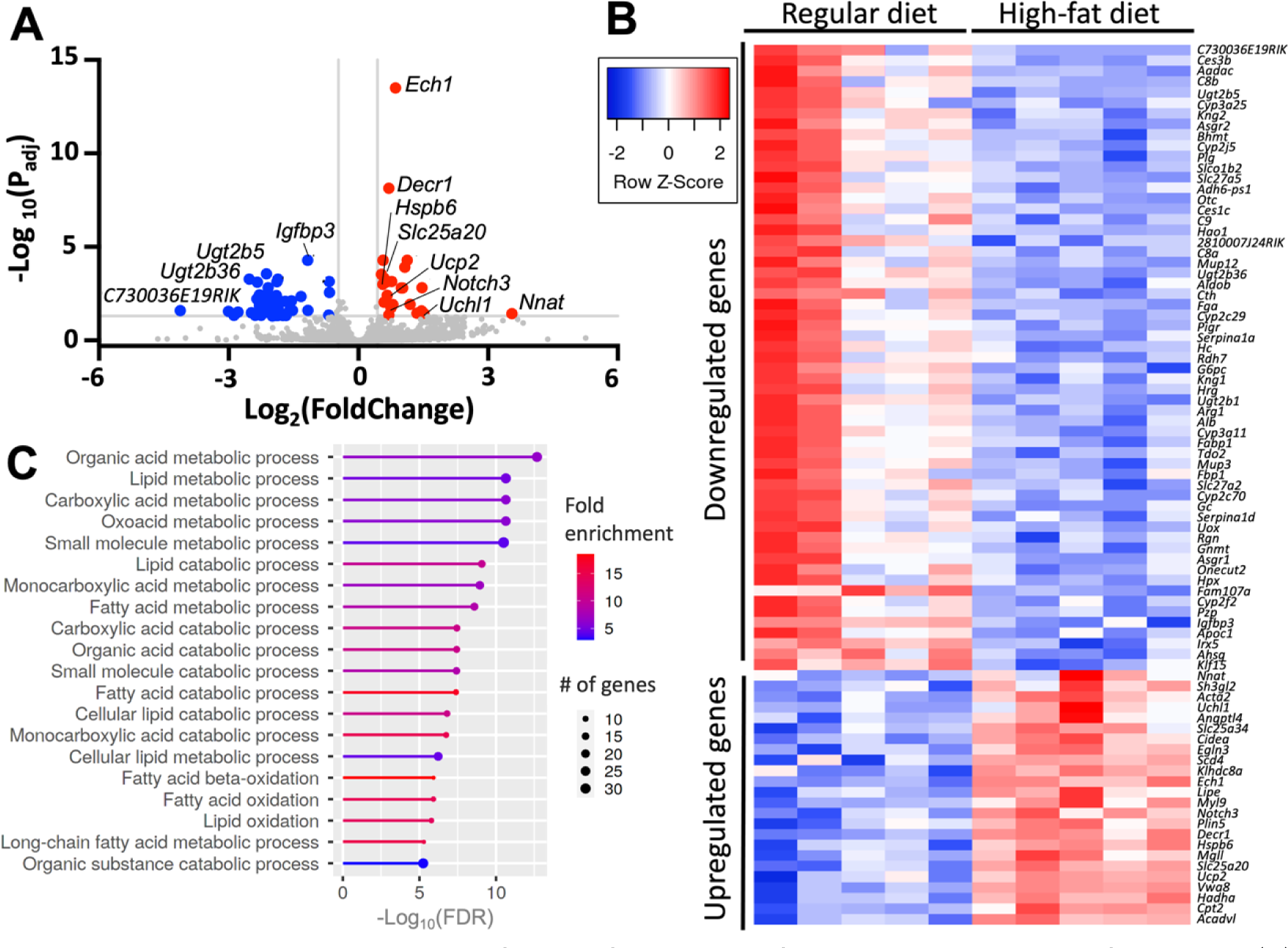
Transcriptomic analysis of atria from mice fed regular and high-fat diets. (A) Volcano plot of log_2_(FoldChange) on the x-axis plotted against -log_10_ adjusted P-value on the y-axis showing upregulated (red) and downregulated (blue) genes induced by diet-induced obesity compared to a regular diet. (B) Pathway analysis showing the gene ontology biological process that is affected by a high-fat diet within the atria and sorted by the adjusted P-value. The color of the bar shows fold change and the circle size indicates the number of differentially expressed genes in each biological process. (C) Heat map of the 59 downregulated and 24 upregulated genes within the atria of mice fed a high-fat diet; N=5 mice per group.

The transcriptomics analysis indicated an upregulation of various genes that are known to contribute to either AF or cardiac remodeling. This included *Uchl1*, which encodes for a ubiquitin hydrolase that is upregulated in myocytes following myocardial infarction [27]; *Slc25a20*, which has been independently associated with rhythm status among patients with AF [28]; and *Upc2*, which influences the susceptibility for Ca^2+^-mediated arrhythmias, modulates myocardial excitation-contraction coupling, and attenuates oxidative stress [29–31]. Conversely, a high-fat diet does not induce changes in gene expression of *Atp2a2* and *Pln*, which encode for SERCA2a and PLN, respectively (Table S3 in the Supplementary Information). We also did not find changes in the gene expression of the Ca^2+^-handling proteins RyR, Ca_v_1.2, NCX1, sarcolipin, and calsequestrin (Table S3 in the Supplementary Information). A high-fat diet also does not affect the gene expression of the Na^+^/K^+^-ATPase (NKA), the NKA-regulating FXYD proteins [32], and the Na_v_1.5 channel (Table S3 in the Supplementary Information). A high-fat diet also induces upregulation of *Hspb6*, a heat-shock protein that protects against remodeling in atrial myocytes upon tachypacing [33], and *Notch3*, a protein that inhibits cardiac fibroblast proliferation [34] (Table S1 of the Supplementary Information). We did not find changes in gene expression of the *Sumo1* and *Sirt1* (Table S3 in the Supplementary Information), two genes that are involved in modulating SERCA2a activity in cardiac cells [35, 36]. Interestingly, *Nnat* is upregulated in mice fed a high-fat diet (Fig. 10 and Table S1 of the Supplementary Information). *Nnat* encodes for neuronatin, a protein that plays a role in whole-body metabolic regulation and has been implicated in obesity [37]. Neuronatin is a SERCA effector that stimulates the SERCA’s ATPase activity [38] and increases SERCA-mediated Ca^2+^ storage [39].

## Discussion

In this study, we used a two-hit model of diet-induced obesity and adrenergic stimulation to investigate the mechanisms by which obesity increases the risk of AF. This model showed expected features of diet-induced obesity, including the increase in body mass and visceral fat and metabolic derangement. Indeed, the model is characterized by glucose intolerance and insulin resistance, in agreement with diet-induced metabolic remodeling [15]. Obesity is a triggering factor for diabetes associated with insulin resistance [40], which increases the risk of arrhythmogenesis and AF [41]. Our model also exhibited increased production of inflammatory cytokines, including TNF-*α* and IL-6. TNF-*α* and IL-6 are inflammatory mediators that have been linked to the pathogenesis of AF [42]. These findings agree with studies showing that obesity promotes and secretion of TNF-*α* and IL-6 by adipose tissue [16, 17].

Histology and Western blot analysis did not reveal either fibrosis or changes in the expression of pro-fibrotic proteins in the atria of mice fed a high-fat diet. These findings contrast with previous studies showing that mice fed a high-fat diet for 10 weeks develop moderate, albeit significant, fibrosis [12, 13]. We used a similar diet to that used in these studies, so we speculate that the diet composition and feeding duration are not variables that help explain the absence of fibrosis in our model. Instead, transcriptomics analysis showed that a high-fat diet induces upregulation of the *Notch3* gene in atria. The Notch signaling pathway is involved in cellular differentiation, proliferation, and apoptosis [43]. Recent studies have shown that *Notch3* inhibits cardiac fibroblast proliferation and fibroblast to myofibroblast transition while promoting cardiac fibroblast apoptosis by modulating the pro-fibrotic RhoA/ROCK/Hif1α signaling pathway [34]. *Notch3* upregulation induced by a high-fat diet may explain the absence of atrial fibrosis observed in our diet-induced obesity mouse model. Future studies are needed to clarify the role of Notch and other anti- and pro-fibrotic signaling pathways in a diet-induced obesity mouse model and their variability with experimental conditions.

Transcriptomics analysis showed that a high-fat diet induces upregulation of *Hspb6*, a heat-shock protein that protects against remodeling in atrial myocytes upon tachypacing [33]. The upregulation of *Hspb6* in our diet-induced obesity model explains why a high-fat diet alone is insufficient to induce AF. Instead, we found that diet-induced obesity and acute sympathetic activation synergize during intracardiac tachypacing to induce AF and that this occurs in the absence of fibrosis [44]. In our model, obesity may promote AF by several mechanisms, including changes in cardiac output, anatomical remodeling of the atria, and systemic hypertension, which can further exacerbate the increase in left ventricular wall stress and left atrium pressure [45]. However, we did not find echocardiographic evidence indicating alterations in left ventricular dynamics, left atrium volume, or differences in systemic blood pressure in our model of diet-induced obesity. These findings agree with studies showing that diet-induced obesity does not affect either left ventricle ejection fraction or systolic blood pressure [46].

At the cellular level, a high-fat diet significantly prolongs the action potential of left atrial myocytes and impairs the effects of *β*-adrenergic stimulation on action potential duration. APD prolongation affects the balance of Ca^2+^ homeostasis, requiring robust compensatory mechanisms [47]. Therefore, we considered early and delayed afterdepolarizations as potential contributors to the action potential prolongation in isolated atrial myocytes. Early afterdepolarizations (EADs) are mainly driven by voltage oscillations in the repolarizing phase of the action potential. However, we did not observe significant changes in I_Ca_ and I_K_ currents in myocytes isolated from mice fed a high-fat diet, suggesting that EADs formation is not the underlying cause for prolonged AP duration [48]. An alternative mechanism for prolongation of the AP is the formation of delayed afterdepolarizations (DADs), which are driven by spontaneous intracellular Ca^2+^ release during diastole [49]. A high-fat diet produces DADs in atrial myocytes, and isoproterenol significantly increases the incidence of DADs induced by diet-induced obesity in mice. These findings are significant because DADs have been related to the initiation of arrhythmias [50], and occur in pathological states including heart failure, diabetes, and ischemic heart disease [51].

Our findings indicate that dysregulation of intracellular Ca^2+^ dynamics contributes to AF in our two-hit model. Therefore, we analyzed the expression of proteins that are involved in Ca^2+^ handling in the atrium. Compared to a regular diet, a high-fat diet does not alter the expression of major proteins involved in Ca^2+^ handling, including RyR, Ca_v_1.2, NCX1, and SERCA2a. We also found that PLN expression is not decreased in response to a high-fat diet. However, a high-fat diet impairs the ability of *β*-adrenergic stimulation to phosphorylate both monomeric and oligomeric forms of PLN. In its unphosphorylated state, PLN inhibits SERCA2a, and PLN phosphorylation reverses this inhibitory effect and reactivates the SERCA2a upon adrenergic activation [52]. The defective PLN phosphorylation found in our model agrees with previous studies that have shown obesity in rats promotes the reduction of PLN phosphorylation [53]. Defective PLN phosphorylation has also been observed in animal models and patient samples of atrial fibrillation [23, 54–56].

Impaired PLN phosphorylation without a reduction in PLN expression suggests that SERCA2a- mediated Ca^2+^ uptake into the sarcoplasmic reticulum is reduced in our diet-induced obesity model. Unexpectedly, we found that Ca^2+^ uptake occurs significantly faster in myocytes from mice fed a high-fat diet *vs* a regular diet. While the Ca^2+^ transient decay reflects the combined activity of SERCA2a and NCX1, the former contributes to ∼90% of Ca^2+^ uptake in mice [57]. Indeed, the Ca^2+^ transient decay in myocytes from obese mice is sensitive to thapsigargin, indicating that the stimulatory effects on Ca^2+^ uptake are primarily mediated by SERCA2a. In atrial cells from obese mice, isoproterenol increases the amplitude of the Ca^2+^ transient but does not affect the relative Ca^2+^ transient decay. These findings and the Western blot analysis indicate that stimulation of Ca^2+^ uptake induced by a high-fat diet is not mediated by PLN phosphorylation. In the absence of *β*-adrenergic stimulation, SERCA2a stimulation exerts negative feedback on Ca^2+^-induced Ca^2+^ release [58], which explains the inability of a high-fat diet to induce AF in the absence of sympathetic stimulation. However, a combination of a high-fat diet and acute sympathetic activation increases sarcoplasmic reticulum Ca^2+^ overloading through SERCA2a activation and decreases sarcoplasmic reticulum Ca^2+^ release threshold, e.g., by RyR phosphorylation [59, 60]. Overall, a high-fat diet primes SERCA2a for activation despite impaired PLN phosphorylation, and the combined effects of a high-fat diet SERCA2a activation and isoproterenol contribute to AF likely through a Ca^2+^-induced Ca^2+^ release gain in atrial myocytes [61, 62]. This mechanism also agrees with a recent study showing that increased intracellular Ca^2+^ mobilization is sensitive to *β*-adrenergic activation, triggering pro-arrhythmia events in ventricular myocytes of mice fed a Western diet [63].

We used transcriptomics analysis to identify gene expression of proteins that may explain stimulation of Ca^2+^ reuptake into the sarcoplasmic reticulum. A high-fat diet does not induce changes in the expression of genes encoding for SERCA2a and PLN. Previous studies have shown SUMO1-dependent stimulation of SERCA2a [35]; however, there were no changes in the expression of the *Sumo1* gene in response to a high-fat diet. Studies have also shown that sirtuin 1-mediated acetylation of SERCA affects the function of this pump [36], but we did not find changes in the expression of the *Sirt1* gene. We found that *Nnat*, which encodes for neuronatin, is upregulated in mice fed a high-fat diet. *Nnat* upregulation agrees with previous studies showing that neuronatin levels correlate with an increase in BMI and body fat mass [64]. Neuronatin is known to activate the SERCA2 isoform [39], thus explaining the paradoxical stimulation of Ca^2+^ uptake in the absence of PLN phosphorylation. Therefore, *Nnat* upregulation induced by a high-fat diet explains the increased Ca^2+^ reuptake into the sarcoplasmic reticulum despite impaired PLN phosphorylation in atrial myocytes.

In summary, we found that diet-induced obesity and acute sympathetic activation synergize during intracardiac tachypacing to induce AF. We used single-cell Ca^2+^ imaging, electrophysiology studies, and Western blot analysis to show paradoxical SERCA2a dysregulation contributes to AF in a model of diet-induced obesity, whereby Ca^2+^ reuptake into the sarcoplasmic reticulum is significantly increased despite a significant reduction in PLN phosphorylation. This finding is significant as it deviates from the current notion that SERCA2a function is generally depressed in cardiac disease [65]. Transcriptomics analysis showed that a high-fat diet promotes upregulation of neuronatin, a modulator of the SERCA pump that is associated with obesity, thus explaining the paradoxical stimulation of Ca^2+^ uptake in our model of diet-induced obesity. Our findings suggest that obesity primes SERCA2a for activation, and adrenergic stimulation facilitates AF conversion likely through a Ca^2+^-induced Ca^2+^ release gain in atrial myocytes.

Clinically, our study suggests that obesity increases the risk of AF upon exposure to acute sympathetic activation. This could occur during the sympathetic nervous system fight-or-flight response, which is mediated by an increase in the spontaneous action potential firing rate of pacemaker cells in the sinoatrial node. These findings have potential translational implications, and pharmacological interventions to mitigate the risk of diet-induced AF may include partial inhibition of SERCA2a [66]. This study suggests a novel mode of SERCA2a dysregulation by which obesity predisposes to atrial fibrillation and has important translational implications to mitigate the risk of this condition in obese patients.

## Materials and Methods

All animal studies were conducted following the guidelines for the Care and Use of Laboratory Animals of the National Institutes of Health and with approval from the University of Michigan Institutional Animal Care and Use Committee.

### High-fat diet-induced obesity mice model

Male C57BL/6 mice were obtained from the Jackson Laboratories (Bar Harbor, ME). Male mice were specifically chosen because the diet-induced obesity model has blunted effects in female mice [67]. Mice were randomly assigned to be fed either a regular chow diet (13.6% fat, 57.5% carbohydrate, 28.9% protein; Research Diet, Labdiet) or a high-fat diet (60% fat, 20% carbohydrate, 20% protein; Research Diet, Labdiet) for 8 or 16 weeks. Body weights were recorded every two weeks. Blood glucose and insulin were measured as previously described [68] and measured at indicated time points. Non-invasive hemodynamics and echocardiography studies were performed at the University of Michigan Physiology and Phenotyping Core. Mice were anesthetized with 2% isoflurane before imaging. We performed B-mode short axis and M-mode imaging and under transthoracic echocardiography using the Vevo 2100 imaging system (FUJIFILM VisualSonics Inc.). A detailed description of the histology studies, ELISA, and Western blot analysis studies performed on mice are described in the Supplementary Information.

### Surface electrocardiographic recording

We performed surface ECG measurements at a one-time point at the end of the feeding period. Mice were anesthetized with 2-3% isoflurane and maintained their body temperature between 36.5-37.5°C using a thermally controlled heating pad. After 10 minutes of stabilization, the ECGs were obtained to establish the baseline (basal) ECG. We then performed an intracardiac surgical procedure to obtain simultaneous surface and intracardiac ECG recordings. Surface ECG analysis was performed as previously described [69].

### Intracardiac recordings

Programmed electrical stimulation was performed as described previously [70] with some modifications and following the guidelines on assessment of arrhythmias in small animals [18]. Mice were anesthetized with 2-3% isoflurane and maintained their body temperature between 36.5-37.5°C using a heating pad. An octapolar catheter (Transonic, Ithaca, NY) was inserted through the jugular vein and advanced into the right atrium and ventricle. Electrical stimulation was performed by using ten atrial bursts pacing before and after intraperitoneal isoproterenol administration (1.5 mg/kg). Atrial stimulation was achieved by rectangular impulses (2 ms) delivered at twice the pacing threshold. A modified S1-S2 burst-stimulation protocol was used to induce atrial arrhythmia: S1 (10 stimuli at cycle length 60ms) was followed by S2 (10 stimuli at cycle length 5ms). This sequence of programmed electrical stimulation was repeated 10 times with a 1-minute delay. We used the PONEMAH (Data Sciences International) software for the acquisition of intracardiac recording data.

### Electrophysiology studies of isolated atrial myocytes

The isolation of atrial myocytes is described in the Supplementary Information. Experiments were carried out using a multi-clamp 700B amplifier (Axon Instruments, Molecular Devices, Union City, CA, USA). Data were acquired and analyzed using the pCLAMP 10 Suite of programs (Axon Instruments,). Borosilicate glass electrodes were pulled with a Brown–Flaming puller (model P- 97), yielding appropriate tip resistances when filled with pipette solution to enable proper voltage control [71, 72]. Action potentials were elicited using square wave pulses 30-50 pA amplitude, 5-8 ms duration, generated by a DS8000 digital stimulator (World Precision Instruments, Sarasota, FL), and recorded at 37°C with pipette solution containing 1 mM MgCl_2_, 1 mM EGTA, 150 mM KCl, 5 mM HEPES, 5 mM phosphocreatine, 4.46 mM K_2_ATP, 2 mM β-Hydroxybutyrate, adjusted to pH 7.2 with KOH. The extracellular solution contained 148 mM NaCl, 0.4 mM NaH_2_PO_4_, 1 mM MgCl_2_, 5.5 mM glucose, 5.4 mM KCl, 1 mM CaCl_2_, 15 mM HEPES, and 1 mM EGTA, pH adjusted to 7.4 with NaOH. RMP, overshoot, action potential amplitude, and APD_25_, APD_50,_ and APD_90_ of repolarization were analyzed using custom-made software, and dV/dt_max_ was calculated using Origin 8.1 (Microcal). We determined the formation of DADs as described previously [73]. Briefly, APs were recorded at 27±5°C with a pipette solution containing 1 mM MgCl_2_, 150 mM KCl, 5 mM HEPES, 5 mM phosphocreatine, 4.46 mM K_2_ATP, 2 mM β-Hydroxybutyrate, with pH adjusted to 7.2 with KOH. A Ca^2+^- and Mg^2+^-containing Hanks’ Balanced Salt Solution (ThermoFisher) was used as an extracellular solution. DADs were recorded before and after treatment with isoproterenol (100 nM). A detailed description of the measurement of I_Ca_ and I_K_ currents is found in the Supplementary Information.

### Single-cell Ca^2+^ imaging

Atrial myocytes were prepared for Ca^2+^ imaging as described in the Supplementary Information. The measurement of intracellular Ca^2+^ transients in atrial myocytes was performed using an Ionoptix recording system. Culture media was removed, and the myocyte-plated coverslips were transferred into Tyrode solution containing the 140 mM NaCl, 4 mM KCl, 1 mM MgCl_2_, 10 mM HEPES, 10 mM Glucose, 1 mM CaCl_2_; the pH was adjusted to 7.4 with NaOH. Cells were loaded with 5 µM Fura2-AM and 2.5 mM probenecid followed by 2 washouts of 10 min each for de-esterification with Tyrode solution. Coverslips were placed in an Ionoptix rapid change stimulation chamber and perfused with Tyrode solution at 37°C; isoproterenol and thapsigargin were tested at a concentration of 200 nM and 1 µM, respectively. A temperature of 37°C was maintained using a mTC3 micro temperature controller. We used an inverted Fluorescence Microscope Nikon ECLIPSE Ti Series and a 40x/1.30 oil Nikon objective for data acquisition. Fura-2 AM excitation wavelengths, 340 nm, and 380 nm were generated using an IonOptix HyperSwitch. Cells were paced (1 Hz and 2 Hz) with the IonOptix Myopacer. A minimum of 10 transients per myocyte were used for ensemble averaging and analyzed using the Ionwizard software (IonOptix LLC, Milton, MA).

### RNA sequencing and differential gene expression

Total RNA was purified from the atria of mice using the Qiagen RNeasy kit (Qiagen, catalog # 74004), and RNA content and quality were determined using the TapeStation System (Agilent). Sequencing was performed at the University of Michigan Advanced Genomics Core (AGC), with Lexogen QuantSeq libraries constructed and subsequently subjected to 101 single-end cycles on the NovaSeq-6000 platform (Illumina). Data were pre-filtered to remove genes with less than 20 counts across all samples. Differential gene expression analysis was performed using DESeq2 [74], using a negative binomial generalized linear model with the following thresholds: linear fold change >1.5 or <-1.5, and a Benjamini-Hochberg FDR <0.05. Genes were annotated with NCBI Entrez GeneIDs [75] and text descriptions. Candidate pathways activated or inhibited comparisons and GO-term enrichments were performed using iPathway Guide (Advaita), iDEP version 0.95 (San Diego State University), and ShinyGO version 0.75 (San Diego State University) [76–78].

### Statistical analysis

All results are presented as mean ± SEM. Normality was determined using the Shapiro-Wilk test. Non-parametric tests were used for data that are not normally distributed. Data were analyzed using the Students T-test or Mann-Whitney U-test for paired experiments, or a two-way analysis of variance (ANOVA) followed by Tukey’s post-hoc test to analyze differences between multiple groups. We used 95% confidence intervals around the differences between the groups for the post-hoc test. Two-sided *p*-values were used, and α-level <0.05 was considered significant.

## Acknowledgments

We thank Laura Martínez Mateu for her assistance in the surface ECG analysis, and Min Zhang for assistance with statistical analysis. This study was funded by the National Institutes of Health grants R01GM120142 and R01HL148068 (to L.M.E.-F), R01AG028082, R35HL155169 and R01AI138347 (to D.R.G.), R00AG068309 (to D.J.T), and American Heart Association grant 20CDA35320040 (to D.P.B). We acknowledge support from the Bioinformatics Core of the University of Michigan Medical School Biomedical Research Core Facilities. This research was supported in part through computational resources and services provided by Advanced Research Computing, a division of Information and Technology Services at the University of Michigan, Ann Arbor.

## Data availability

All data are available in the main text and supplementary information.

