## Supplementary information for "Paradoxical SERCA dysregulation contributes to atrial fibrillation in a model of diet-induced obesity"

**This PDF file includes:**

Extended methods
Tables S1 to S3

**Extended methods**

***Histology***. Blood was removed by right ventricular puncture and the vasculature was perfused with ice-cold PBS. The heart was harvested, and isolated RA and LA were fixed in 4% paraformaldehyde, embedded in paraffin, and cut into 5 mm sections. Masson trichrome staining was used to evaluate interstitial fibrosis. Hematoxylin and eosin staining were used to evaluate fat accumulation. Paraffin-embedded sections of the fat pads were deparaffinized and rehydrated. After blocking, sections (6µm each) were incubated at room temperature for 2 hours with Mac2 (sc-81728; Santa Cruz Biotechnology Inc.). Mac2 slides were counterstained with hematoxylin and cover-slipped. Images were captured with an Olympus LC30 camera mounted on an Olympus CX41 microscope. For the $Mac2^+$ area, all images were obtained with the same light source at the same time. The $Mac2^+$ area was determined using the threshold function in ImageJ (NIH) and normalized to the total region of interest area. Results are reported as the percentage of adipose area. Sectioning and staining were performed by the In Vivo Animal Core Laboratory technicians at the Unit for Laboratory Animal Medicine, University of Michigan. Technicians in this laboratory were blinded to experimental identity.

***ELISA***. Left perigonadal fat pads were isolated from HFD-fed or LFD-fed mice and incubated for 24 hours in RPMI 1640 plus 10% fetal bovine serum with 1% penicillin-streptomycin solution (P4333; Sigma-Aldrich). Conditioned, culture supernatants were collected and stored at -80°C. ELISAS for IL-6 (ThermoFisher # KMC0061), TNFα (ThermoFisher # BMS607-3), and galectin-3 (R&D Systems, # DY1197) were performed according to the manufacturer's instructions and normalized to fat pad weight.

***Western blot analysis***. Hearts were harvested in cold 4°C PBS, and then both right and left atria were separated from the ventricles, homogenized in 100 µL lysis buffer (ThermoFisher Scientific, catalog # 78510) with 1% protease inhibitor cocktail (Sigma, catalog # P8340) and 1% phosphatase inhibitor cocktail (Sigma, catalog # P5726). Atrial tissue lysates in b-mercaptoethanol 4X sample buffer were loaded into 4-12% NuPage precast gels (ThermoFisher, Waltham, MA USA) and electrophoresis was carried out. The SDS-PAGE resolved proteins were transferred to iBlot® stacks with regular PVDF membranes using the Life Technologies iBlot2 system. Non-specific binding sites were blocked with 5% bovine serum albumin (BSA) in PBS-T (in mM, 3 $KH_2PO_4$, 10 $Na_2HPO_4$, 150 NaCl, 0.1% Tween 20, pH 7.2-7.4) for 30 min at room temperature. Membranes were then incubated with specific primary antibodies diluted in 5% BSA in PBS-T overnight at 4°C. After washing 3 times for 10 min, membranes were incubated with horseradish peroxidase-conjugated secondary antibodies diluted in 5% BSA in PBS-T for 1 h. After washing 3 times for 10 minutes, protein-antibody reactions were detected by Supersignal chemiluminescence (Pierce Biotechnology Inc, Rockford, IL, USA) and imaged using Image Lab software 5 (Bio-Rad). Densities of proteins were measured using the Image Lab software version 5.

***Isolation of cardiac myocytes***. Mice cardiac myocytes were isolated using a modified Langendorff technique. Briefly, mice were intraperitoneal injected with 0.1 mL heparin (1000 IU/mL) 20 min before heart excision. Animals were anesthetized with 2% isoflurane and the heart was removed quickly from the chest and retrogradely perfused through the aorta at a constant flow (4 mL/min) at 37 $^0$C for 4 min with a $Ca^{2+}$-free buffer containing 113 mM NaCl, 4.7 mM KCl, 1.2 mM $MgSO_4$, 0.6 mM $Na_2HPO_4$, 0.6 mM $KH_2PO4$, 10 mM $KHCO_3$, 12 mM $NaHCO_3$, 10 mM HEPES, 10 mM 2,3-butanedione monoxime (BDM, sigma) 30 mM taurine, and 5.5 mM glucose. All solutions were filtered (0.2 mm filter) and equilibrated with $O_2$ (100%) for at least 20 min before use. Enzymatic digestion was initiated by adding collagenase type II (773.4 U/mL; Worthington), trypsin (0.14 mg/mL), and $CaCl_2$ (12.5 mM) to the perfusion solution. After 5-6 min of digestion, the atrial tissue was removed and placed in a well of a 24-well plate with enzymatic solution, stirred gently with a magnetic bar for 5 minutes. After this time the isolated atrial myocytes were collected, and the remaining tissue was transferred to another well containing enzymatic solution. This was repeated until the tissue was completely dissociated.

***Measurement of $I_{Ca}$ and $I_K$ currents***. For $I_{Ca}$ measurements, we used the following pipette filing solution: 120 mM CsCl, 20 mM TEA-Cl, 1 mM $MgCl_2$, 5 mM MgATP, 0.2 mM $Na_2GTP$, 10 mM HEPES, and 10 mM EGTA, pH adjusted to 7.2 with CsOH. The external solution contained 137 mM NaCl, 5.4 mM CsCl, 1 mM $MgCl_2$, 1 mM $CaCl_2$, 10 mM HEPES, 10 mM glucose, 2 mM 4-aminopyridine (4-AP), and 10 mM tetrodotoxin (TTX); pH was adjusted to 7.4 with NaOH. Voltage dependence of peak $I_{CaL}$ was measured by holding at -50 mV; 300 ms voltage steps were applied from -40 to +60 mV in 5 mV increments. The interval between voltage steps was 350 ms. For $I_K$ measurements, we used the following pipette filling solution: 150 mM KCl, 1 $MgCl_2$, mM 5 EGTA, 5 mM HEPES, 5 mM phosphocreatine, 4.4 mM $K_2ATP$, 2 mM β-hydroxybutyric acid, pH adjusted to 7.2 with KOH. The external solution contained 148 mM NaCl, 0.4 mM $NaH_2PO_4$, 1 mM $MgCl_2$, 5.5 mM glucose, 5.4 mM KCl, 1 mM $CaCl_2$, and 15 mM HEPES; the pH was adjusted to 7.4 with NaOH. To measure $I_K$ currents, we used 10 mM TTX and 5 mM nifedipine to block the fast voltage-gated $Na^+$ channels and the L-type $Ca^{2+}$ currents, respectively. $K^+$ currents were recorded using 5-second depolarizing pulses to potentials between -120 mV to +50 mV from a holding of -70 mV. Voltage steps were in steps of 10 mV at 10-second intervals. Cells were constantly perfused with the appropriate external solution. All membrane currents were examined as current densities by normalizing to cell capacitance. For acute adrenergic stimulation, we used 10 μM isoproterenol.

***Preparation of atrial myocytes for $Ca^{2+}$ imaging studies***. After myocyte dissociation, calcium reintroduction was gradually performed to a final concentration of 1 mM. The myocytes were allowed to equilibrate at room temperature for 10 min and transferred to a plating media solution with the following composition: culture media (see below) supplemented with 5% Fetal bovine serum (Sigma, F2442) and 25 μM S-(-) Blebbistatin (Sigma, B0560). Myocytes were counted using a hemocytometer, and approximately 20,000 cells were plated on previously 22x22 mm glass coverslips coated with laminin (40 μg/mL) using 6-well plates. The cells were allowed to attach for 40 min at room temperature. After cell attachment, the plating media was replaced with culture media with the following composition: 150 mL MEM +Hank's salts (Gibco, 500 mL 11575-032), 0.1% ITS liquid media supplement (100x, Sigma, I3146), 10 U/mL Penicillin/ 10 μg/mL Streptomycin (Invitrogen, 15070-063), 25 mM NaHCO3 (Sigma, S8875), 15 mM HEPES (Sigma H7006), 5 mM butanedione monoxime (BDM, sigma), and the cells were allowed to equilibrate for 20 min at room temperature.

**Table S1.** Results from the transcriptomics analysis showing genes that are upregulated in atria of mice fed a high-fat diet.

| Gene ID | Description | $\log_2$(FoldChange) | $P_{adj}$ |
|---|---|---|---|
| *Nnat* | Neuronatin | 3.9358749 | 3.75E-02 |
| *Sh3gl2* | SH3-domain GRB2-like 2 | 1.6766716 | 3.67E-02 |
| *Acta2* | Actin $\alpha$-2 | 1.6285951 | 1.51E-03 |
| *Uchl1* | Ubiquitin carboxy-terminal hydrolase L1 | 1.6152538 | 2.66E-02 |
| *Angptl4* | Angiopoietin-like 4 | 1.5013384 | 3.55E-02 |
| *Slc25a34* | Solute carrier family 25, member 34 | 1.3218376 | 1.17E-02 |
| *Cidea* | Cell death-inducing DNA fragmentation factor | 1.2472558 | 5.29E-05 |
| *Egln3* | egl-9 family hypoxia-inducible factor 3 | 1.1866723 | 1.22E-04 |
| *Scd4* | Stearoyl-coenzyme A desaturase 4 | 1.1364512 | 1.60E-03 |
| *Klhdc8a* | Kelch domain-containing 8A | 1.0969114 | 1.60E-03 |
| *Ech1* | Enoyl coenzyme A hydratase 1 | 0.9524622 | 3.45E-14 |
| *Lipe* | Lipase, hormone-sensitive | 0.8801781 | 1.22E-02 |
| *Myl9* | Myosin, light polypeptide 9 | 0.854425 | 7.00E-04 |
| *Notch3* | Notch receptor 3 | 0.8333349 | 1.78E-02 |
| *Plin5* | Perilipin 5 | 0.7815313 | 4.34E-02 |
| *Decr1* | Mitochondrial 2,4-dienoyl CoA reductase 1 | 0.7790149 | 7.42E-09 |
| *Hspb6* | Heat shock protein, $\alpha$-crystallin-related, B6 | 0.7380947 | 7.27E-04 |
| *Mgll* | Monoglyceride lipase | 0.7357712 | 3.92E-03 |
| *Slc25a20* | Solute carrier family member 20 | 0.6602545 | 4.78E-04 |
| *Ucp2* | Uncoupling protein 2 | 0.6542212 | 9.06E-03 |
| *Vwa8* | Von Willebrand Factor A Domain Containing 8 | 0.635321 | 5.29E-05 |
| *Hadha* | $\alpha$-subunit, mitochondrial trifunctional protein | 0.6298784 | 5.29E-05 |
| *Cpt2* | Carnitine palmitoyltransferase 2 | 0.6259808 | 9.94E-04 |
| *Acadvl* | Acyl-Coenzyme A dehydrogenase, long chain | 0.5910815 | 2.94E-04 |

**Table S2.** Results from the transcriptomics analysis showing genes that are downregulated in atria of mice fed a high-fat diet.

| Gene ID | Description | $\log_2$(FoldChange) | $P_{adj}$ |
|---|---|---|---|
| *Hao1* | Hydroxyacid oxidase 1, liver | -4.56117892 | 2.59E-02 |
| *2810007J24RIK* | Non-coding RNA lncLSTR | -3.32859513 | 2.77E-02 |
| *C8a* | Complement component 8,α-polypeptide | -3.18667035 | 4.61E-02 |
| *Mup12* | Major urinary protein 12 | -3.08277672 | 3.00E-02 |
| *Ugt2b36* | UDP glucuronosyltransferase 2, PP 36 | -2.79395868 | 5.35E-04 |
| *Aldob* | Aldolase B, fructose-bisphosphate | -2.74285967 | 3.29E-02 |
| *Cth* | Cystathionase (cystathionine γ-lyase) | -2.64470838 | 4.61E-02 |
| *Fga* | Fibrinogen α-chain | -2.60214385 | 6.01E-03 |
| *Cyp2c29* | Cytochrome P450, family 2, PP29 | -2.58927548 | 7.70E-04 |
| *Pigr* | Polymeric immunoglobulin receptor | -2.53607478 | 1.03E-02 |
| *Serpina1a* | Serine/Cysteine peptidase inhibitor 1A | -2.53323819 | 3.64E-03 |
| *Hc* | Hemolytic complement | -2.53075027 | 1.42E-02 |
| *Rdh7* | Retinol dehydrogenase 7 | -2.49011343 | 4.61E-02 |
| *G6pc* | Glucose-6-phosphatase | -2.48081814 | 7.64E-03 |
| *Kng1* | Kininogen 1 | -2.47785248 | 2.66E-02 |
| *Hrg* | Histidine-rich glycoprotein | -2.44995622 | 1.97E-02 |
| *Ugt2b1* | UDP glucuronosyltransferase 2, PP B1 | -2.43224654 | 3.30E-02 |
| *Arg1* | Arginase | -2.41928113 | 2.59E-02 |
| *Alb* | Albumin | -2.39526486 | 4.21E-03 |
| *Cyp3a11* | Cytochrome P450, family 3, PP11 | -2.39243285 | 3.10E-02 |
| *Fabp1* | Fatty acid binding protein 1 | -2.35790185 | 1.21E-02 |
| *Tdo2* | Tryptophan 2,3-dioxygenase | -2.35455506 | 2.78E-04 |
| *Mup3* | Major urinary protein 3 | -2.33196502 | 1.10E-02 |
| *Fbp1* | Fructose bisphosphatase 1 | -2.32152002 | 3.92E-03 |
| *Slc27a2* | Solute carrier family 27 member 2 | -2.31684202 | 2.89E-03 |
| *Cyp2c70* | Cytochrome P450, family 2, PP70 | -2.06324976 | 4.74E-02 |
| *Gc* | GC Vitamin D Binding Protein | -2.06264149 | 5.41E-04 |
| *Serpina1d* | Serine/Cysteine peptidase inhibitor 1D | -2.0567446 | 1.99E-02 |
| *Uox* | Urate oxidase | -2.04854547 | 2.65E-02 |
| *Rgn* | Regucalcin | -1.93700546 | 4.71E-02 |
| *Gnmt* | Glycine N-methyltransferase | -1.87548375 | 8.05E-03 |
| *Asgr1* | Asialoglycoprotein receptor 1 | -1.85369659 | 4.71E-02 |
| *Onecut2* | One-cut domain, family member 2 | -1.7671305 | 2.66E-02 |
| *Hpx* | Hemopexin | -1.76009049 | 2.51E-02 |
| *Fam107a* | Family with seq. similarity 107, member A | -1.71388062 | 7.51E-03 |
| *Cyp2f2* | Cytochrome P450, family 2, subfamily F, PP2 | -1.68980162 | 2.49E-02 |
| *Pzp* | PZP, α-2-macroglobulin like | -1.46863366 | 4.52E-03 |
| *Igfbp3* | Insulin-like growth factor binding protein 3 | -1.301052 | 5.29E-05 |
| *Apoc1* | Apolipoprotein C-I | -1.29237357 | 2.44E-02 |
| *Irx5* | Iroquois homeobox 5 | -0.76118912 | 4.44E-02 |
| *Ahsg* | α-2-HS-glycoprotein | -0.74651702 | 7.00E-04 |
| *Klf15* | Kruppel-like factor 15 | -0.74499343 | 2.74E-03 |

**Table S3.** Summary of gene expression of calcium-handling, sodium-transporting, and SERCA2a-regulating proteins in atria of mice fed a high-fat diet.

| Gene ID | Description | $log_2$(FoldChange) | $P_{adj}$ |
|---|---|---|---|
| *Atp2a2* | Sarcoplasmic reticulum $Ca^{2+}$-ATPase isoform 2 (SERCA2a) | -0.09544436 | 0.011429456 |
| *Pln* | Phospholamban | -0.46219364 | 1.786313278 |
| *Ryr2* | Cardiac ryanodine receptor 2 (RyR2) | 0.1153773 | 0.014622124 |
| *Cacna1c* | $Ca_v$1.2 L-type voltage-gated calcium channel | -0.07287843 | 0.004578043 |
| *Slc8a1* | $Na^+/Ca^{2+}$ exchanger protein 1 | 0.2411005 | 0.147312776 |
| *Casq2* | Calsequestrin 2 | 0.2015695 | 0.248789215 |
| *Sln* | Sarcolipin | -0.24274848 | 0.398687932 |
| *Atp1a1* | $Na^+/K^+$ ATPase transporting subunit alpha 1 (NKA-$\alpha$1) | 0.2054246 | 0.045051404 |
| *Atp1a2* | $Na^+/K^+$ ATPase transporting subunit alpha 2 (NKA-$\alpha$2) | 0.01571423 | 0.000645076 |
| *Fxyd1* | Phospholemman | 0.1064471 | 0.009333479 |
| *Fxyd2* | FXYD Domain-Containing Ion Transport Regulator 2 | 1.0412618 | 0.072278579 |
| *Fxyd5* | Dysadherin | 0.4837579 | 0.446589599 |
| *Fxyd6* | FXYD Domain-Containing Ion Transport Regulator 6 | 0.4823985 | 0.860362129 |
| *Fxyd7* | FXYD Domain-Containing Ion Transport Regulator 7 | -0.13341468 | 0.000645076 |
| *Scn5a* | Sodium Voltage-Gated Channel Alpha Subunit 5 ($Na_v$1.5) | 0.2409932 | 0.117074975 |
| *Sirt1* | Sirtuin 1 | -0.224188348 | 0.854507362 |
| *Sumo1* | Small Ubiquitin Like Modifier 1 | 0.078081544 | 0.989338099 |
